# An enveloped virus-like particle vaccine expressing a stabilized prefusion form of the SARS-CoV-2 spike protein elicits potent immunity after a single dose

**DOI:** 10.1101/2021.04.28.441832

**Authors:** Anne-Catherine Fluckiger, Barthelemy Ontsouka, Jasminka Bozic, Abebaw Diress, Tanvir Ahmed, Tamara Berthoud, Anh Tran, Diane Duque, Mingmin Liao, Michael McCluskie, Francisco Diaz-Mitoma, David E. Anderson, Catalina Soare

## Abstract

Development of efficacious single dose vaccines would substantially aid efforts to stop the uncontrolled spread of the COVID-19 pandemic. We evaluated enveloped virus-like particles (eVLPs) expressing various forms of the SARS-CoV-2 spike protein and several adjuvants in an effort to identify a COVID-19 vaccine candidate efficacious after a single dose. The eVLPs expressing a modified prefusion form of SARS-CoV-2 spike protein were selected as they induced the highest antibody binding titers and neutralizing activity after a single injection in mice. Formulation of SARS-CoV-2 S eVLPs with aluminum phosphate resulted in balanced induction of IgG2 and IgG1 isotypes and antibody binding and neutralization titers were undiminished for more than 3 months after a single immunization. A single dose of this candidate, VBI-2902a (prefusion S eVLPs formulated with aluminum phosphate), protected Syrian golden hamsters from challenge with SARS-CoV-2 and supports the on-going clinical evaluation of VBI-2902a as a potential single dose vaccine against COVID-19.

**Highlights:**

- VBI-2902a is a VLP-based vaccine candidate against SARS-COV-2
- VBI-2902a contains VLPs pseudotyped with a modified prefusion SARS-COV-2 S in Alum.
- VBI-2902a induces robust neutralization antibody response against SARS-COV-2 S
- VBI-2902a protects hamsters from SARS-CoV-2 induced lung inflammation
- A single dose of VBI-2902a provides protective benefit in hamsters

## 1. Introduction

SARS-CoV-2 has been circulating worldwide for more than a year with no significant sign of natural exhaustion, in contrast to the previous SARS and MERS epidemics in 2003 and 2012 respectively, which faded despite absence of a vaccine or specific antiviral treatments. In contrast to the SARS epidemic, the COVID-19 pandemic is associated with increasing morbidity, mortality, and mutagenic potential as more people are infected at an increasing rate [1]. The possibility of waning immunity and isolated cases of re-infection after a period of convalescence have been reported that have prompted questions about correlates of protection and the efficacy of natural immunity [2-4]. Unprecedented efforts and measures have been undertaken to rapidly provide prophylactic vaccines that could decrease the rate of infection and prevent severe health complications.

The SARS-CoV-2 spike (S) protein was identified as a major target for neutralizing antibodies (nAb) due to its crucial role in mediating virus entry and its homology to S proteins from SARS, MERS and other coronaviruses (CoV) for which nAb had similarly been demonstrated [5,6]. CoV S proteins resemble typical of class I viral proteins. They are constituted of 2 functional subunits, S1, containing the receptor binding domain (RBD) and S2, containing the fusion entry domain. Binding of the RBD to the host cell receptor induces conformational changes resulting in activation of the protease cleavage site upstream of the fusion domain followed by release and activation of the S2 fusogenic domain [7]. Unlike SARS-CoV and other CoV from the same clade, SARS-CoV-2 S contains a furin cleavage site located at the boundary of S1 and S2 [6,8](Walls, Cell 2020: Coutard, Antiviral Res. 2020) enabling rapid processing of the S protein during biosynthesis in host cells.

The CoV S proteins are expressed at the viral surface as metastable prefusion trimers that undergo conformational changes [6-7]. Studies of class I viral fusion proteins resulted in the design of stabilized prefusion forms resistant to protease cleavage that could increase expression yield and elicit potent neutralization responses in mice [9-10]. Wrapp et al. [11] described a similar strategy whereby 2 consecutive prolines in the S2 subdomain between heptad repeat 1 and the central helix are substituted with the addition of a C-terminus foldon trimerization domain. The result was a SARS-COV-2 stabilized S-2P antigen. Vaccine candidates containing SARS-CoV-2 S-2P have demonstrated potent induction of nAb responses in laboratory animals [12-13] and humans [14-15].

Virus-like particles (VLPs) are attractive vaccine candidates to generate nAb responses. Structurally, they resemble the wild-type virus from which they are derived, but are much safer because they lack genetic material and therefore the ability to replicate [16]. VLPs enable repeating, array-like presentation of antigens which is a preferred means of activating B cells and eliciting high affinity antibodies [17]. Indeed, VLP expression of a B cell antigen improved neutralizing titers over 10-fold relative to immunization with the same amount of recombinant protein [18]. Accordingly, the use of VLPs as a vaccine modality may expand higher affinity B cell repertoires relative to recombinant protein or DNA/mRNA-based modalities.

In the present study murine leukemia virus (MLV)-based enveloped virus-like particles (eVLPs) [18,19] were used to produce vaccine candidates expressing various forms of SARS-CoV-2 S. We demonstrate that a modified prefusion form of S containing the ectodomain of SARS-CoV-2 S fused with the transmembrane cytoplasmic terminal domain of VSV-G enabled high yields and density of S expression on MLV-Gag eVLPs and induced robust nAb responses exceeding those observed with SARS-CoV-2 convalescent sera after a single dose when adjuvanted with Alum phosphate (VBI-2902a). This candidate, VBI-2902a, was safe and highly efficacious in a hamster challenge model and offers the potential for use as a single dose vaccine. More generally, these data demonstrate the high potency of antigen expression by eVLPs.

## 2. Materials and Methods

### 2.1. COVID-19 human sera

Plasma samples were purchased from Biomex GmbH (Heidelberg, Germany). Samples were collected under consent at donation centers in Heildelberg or Munich, from 30 individual who recovered from moderate SARS-CoV-2 infection with no need for hospitalization or heavy treatment. Subjects were aged 26 to 61 years old. Sera were collected at 26 to 72 days post time of infection. One 61 years old woman was asymptomatic but all others experienced multiples symptoms including fever, headache, anosmia, coughing, difficulty to breath, tiredness and muscle pain.

### 2.2. Plasmids, eVLPs production and adjuvant formulation

All sequences coding for the full length and modified S proteins from SARS-CoV-2 were codon optimized prior to synthesis and subcloned into a proprietary modified phCMV plasmid at Genscript (Piscataway, NJ). The proprietary HEK293SF-3F6 GMP compliant cells were provided by the National Research Council (NRC, Montreal, Canada) and grown in serum-free chemically defined medium [20]. eVLPs were produced using transient polyethylenimine transfection in 293SF-3F6 by co-transfection of one plasmid coding for the spike protein with a phCMV plasmid encoding minimal cDNA sequence of murine leukemia virus (MLV) Gag corresponding to the full length Gag deprived of its C-terminal Polymerase sequence as described elsewhere [18]. Control called “empty” eVLPs or Gag eVLPs were produced by exclusive transfection of the Gag plasmid. Cell culture harvests containing eVLPs were processed using a proprietary purification steps that consists of clarification, tangential flow filtration, benzonase® treatment, diafiltration and ultracentrifugation using sucrose cushion. The final product was sterile filtered using 0.2 µm membrane prior to preparation of vaccine. Depending on the different pre-clinical mouse studies, SARS-CoV-2 eVLPs vaccines were formulated with different adjuvant system including Alum (Adjuphos®), AS03, MF59, and AS04. Adjuvants AS03, ASO4 and its VBI modified version, AddavaxTM (MF59), 2% AdjuPhos® (aluminium phosphate refered as Alum) were purchased from Invivogen.

### 2.3. Western blot analysis of eVLPs content

The expression of SARS-CoV-2 S protein in eVLP preparations was analyzed by western blotting as described previously [18] using rabbit polyclonal Ab (pAb) anti-spike protein of SARS-CoV-2 (Immune Technology Corp) followed by detection with goat anti-rabbit IgG-Fc horseradish peroxydase-conjugated (Bethyl). Alternatively, human sera from COVID-19 convalescent subjects was used as primary antibody followed by detection with goat anti-human IgG heavy and light chain HRP-conjugated (Bethyl). Precision Protein Streptactin HRP conjugate (Bio-Rad) was used as molecular weight ladder standard. Recombinant SARS-COV-2 S(S1+S2) unmodified protein (Mybiosource) or SARS-CoV-2 stabilized prefusion S protein (National Research Council of Canada -NRC) were used as controls.

### 2.4. Mouse immunization study

Six- to 8-week-old female C57BL/6 mice were purchased from Jackson Laboratory (ME, USA). The animals were allowed to acclimatize for a period of at least 7 days before any procedures were performed. The animal studies were conducted under ethics protocols approved by the National Research Council of Canada Animal Care Committee. The animals were maintained in a controlled environment in accordance with the “Guide for the Care and Use of Laboratory Animals” at the NRC Animal Research facility (Institute for Biological Sciences, Ottawa, Canada). Mice were randomly assigned to experimental groups and received intraperitoneal (IP) injections with 0.5 mL of different adjuvanted SARS-CoV-2 immunogens. Blood was collected on day -1 (pretreatment) and day 14 after each injection. All mice from each group were sacrificed 14 days after the last immunization for humoral immunity assessment and or 6 days after the second immunity, where spleens were collected for cellular immunity assessment.

### 2.5. Hamster challenge study

Syrian golden hamsters (males, 5-6 weeks old) were purchased from Charles River Laboratories (Saint-Constant, Quebec, Canada). The study was conducted under approval of the CCAC committee at the Vaccine and Infectious Disease Organization (VIDO) International Vaccine Centre (Saskatchewan,Canada). Animals were randomly assigned to each experimental groups (A, B) (n=12/group) in two independent experiments (Regimen II and Regimen I). Groups A placebo received 0.9%-saline buffer, Groups B received VBI-2902a. Each dose of VBI-2902a contained 1µg of SPG and 125 µg of Alum. Injection was performed by intramuscular (IM) route at one side of the thighs in a 100 µL volume. The schedule for immunization, challenge and sample collection was depicted on Fig. 6a. All animals were challenged intranasally via both nares with 50 μL/nare containing 1×10^5^ TCID50 of SARS-CoV-2/Canada/ON/VIDO-01/2020/Vero’76 (Seq. available at GISAID EPI_ISL_413015) strain per animal. Body weights and body temperature were measured at immunization for 3 days and daily from the challenge day. General health conditions were observed daily through the entire study period. Blood samples and nasal washes were collected as indicated on Fig. 6a. Half of the animals (6/group) were euthanized at 3 days post-infection (dpi), and the remaining animals were euthanized at 14 dpi. The challenge experiments were performed in the animal biosafety level 3 (ABSL3) laboratory at VIDO (Saskatchewan, Canada).

### 2.6. Antibody binding titers

Anti-SARS-CoV-2 specific IgG binding titers in mouse sera were measured by standard ELISA procedure described elsewhere [18], using recombinant SARS-CoV-2 S (S1+S2) protein (Sinobiological). For total IgG binding titers, detection was performed using a goat anti-mouse IgG-Fc HRP (Bethyl) for mouse serum, or goat anti-human IgG heavy and light chain HRP-conjugated (Bethyl) for human serum. HRP-conjugated Goat anti-mouse IgG1 and HRP-conjugated goat anti-mouse IgG2b HRP (Bethyl) were used for the detection of isotype subtype. Determination of Ab binding titers to the RBD was performed using SARS-COV-2 RDB recombinant protein (Sinobiological). Detection was completed by adding 3,3’,5,5’-tetramethylbenzidine (TMB) substrate solution, and the reaction stopped by adding liquid stop solution for TMB substrate. Absorbance was read at 450 nm in an ELISA microwell plate reader. Data fitting and analysis were performed with SoftMaxPro 5, using a four-parameter fitting algorithm. Ab binding titers in hamster sera were determined with ELISA method. Plates were coated with spike S1+S2 Ag (Sinobiological). The coating concentration was 0.1 ug/mL. Plates were blocked with 5% non-fat skim milk powder in PBS containing 0.05% Tween 20. Fourfold dilutions of serum were used. Goat anti-Hamster IgG HRP from ThermoFisher (PA1-29626) was used as the secondary antibody at 1:7000. Plates were developed with OPD peroxidase substrate (0.5 mg/ml) (Thermo Scientific Pierce). The reaction was stopped with 2.5 M sulfuric acid and absorbance was measured at 490 nm. Throughout the assay, plates were washed with PBS containing 0.05% Tween 20. The assay was performed in duplicate. The titres were reported as the end point of the dilutions.

### 2.7. Virus neutralization assays

Neutralizing activity in mouse serum samples was measured by standard plaque reduction neutralization test (PRNT) on Vero cells at the NRC (Ottawa, Canada) using 100 PFU of SARS-CoV-2/Canada/ON/VIDO-01/2020. Results were represented as PRNT90, PRNT80, or PRNT50 end point titer, corresponding to the lowest dilution inhibiting respectively 90% or 80% or 50% of plaque formation in Vero cell culture.

Virus neutralization assays against the challenge SARS-CoV-2 virus were performed at VIDO, Saskatchewan on the hamster serum samples collected at pre-challenge and at the end day; 3 days post-challenge or 14 days post-challenge. The study was conducted using the cell line Vero E6. The serum samples were heat-inactivated for 30 min at 56°C. The serum samples were initially diluted 1:10 and then serially diluted (2-fold serial dilutions). The virus was diluted in medium for a final concentration of 3×10^2^ TCID50/mL. Initially 60 μL of the virus solution was mixed with 60 μL serially diluted serum samples. The mixture was incubated for 1hr at 37°C, with 5% CO2. The pre-incubated virus-serum mixtures (100 μL/well) were transferred to the wells of the 96-well flat-bottom plates containing 90% confluent pre-seeded VeroE6 cells. The plates were incubated at 37°C, with 5% CO2 for 5 days. The plates were observed using a microscope on day 1 post-infection for contamination and on days 3 and 5 post-infection for cytopathic effect (CPE). The serum dilution factor for the last well with no CPE at 5 dpi was defined as the serum neutralization titer. The initial serum dilution factor was 1:20.

### 2.8. RNA extraction and purification

RNA was extracted using QIAamp Viral RNA Mini Kit (Qiagen). Briefly, 140 μL of hamster nasal wash was added into 560 μL viral lysis buffer (Buffer AVL). The mixture was incubated at room temperature for 10 min. After brief centrifugation, the solution was transferred to a fresh tube containing 600 μL of 100% ethanol, and the tube was incubated at room temperature for 10 min. RNA was then purified using QiaAmp Viral RNA Mini Kit and eluted with 60 μL of RNase Free water containing 0.04% sodium azide (elution buffer AVE). Extraction of RNA from lung lobes and nasal turbinates was completed using approximately 100 μg of tissue. The tissues were homogenized in 600 μL of lysis buffer (RLT Qiagen) with a sterile stainless steel bead in the TissueLyserII (Qiagen) for 6 min, at 30 Hz. The solution was centrifuged at 5000 x g for 5 min. Supernatant was transferred to a fresh tube containing 600 μL of 70% ethanol, and the tube was incubated at room temperature for 10 min. Viral RNA was then purified using Qiagen Rneasy Mini Kit (Cat No /ID: 74106) and eluted with 50 μL elution buffer.

### 2.9. Viral qRT-PCR reaction

The qRT-PCR assays were performed on RNA from samples of nasal washes, lung tissues and nasal turbinates using SARS-CoV-2 specific primers targeting the E gene (Fwd, *ACAGGTACGT-TAATAGTTAATAGCGT;* Rev, *ATATTGCAGCAGTACGCACACA*) and labelled probe, *ACACTA-GCCATCCTTACTGCGCTTCG*. The primers have an annealing temperature of approximately 60°C. Qiagen Quantifast RT-PCR Probe kits were used for qRT-PCR. The qRT-PCR results were expressed in RNA copy number per reaction. This was done by producing a standard curve with RNA extracted from a sample of SARS-CoV-2 which was cloned to determine exact copy number of the gene of interest. The Ct values for individual samples were used with the standard curve to determine the copy number in each sample. The qRT-PCR reactions were performed using the OneStepPlus (Applied Biosystems) machine. The program was set at: Reverse transcription (RT) 10 min at 50°C; Inactivation 5 min at 95°C; and then 40 cycles of denaturation for 10 sec at 95°C and annealing/extension for 30 sec at 60°C.

### 2.10. IFN-γ Ex-vivo ELISPOT

IFN-γ ELISPOT analyses to measure Th1 T cell responses were performed as follows. One day before the spleens were removed, ELISpot plates (Millipore) were coated with IFN-γ capture antibody at a concentration of 15 µg/mL (Mabtech). The following day, mice were sacrificed and spleens were removed. Spleens from individual mice were processed to produce single cell suspensions. Erythrocytes were lysed using a commercially available RBC lysis buffer (BioLegend). Fifty microliters containing 2×10^6^ splenocytes were then to each well of a pre-blocked ELISPOT plate. Then, fifty microliters of stimulant pepmixes (JPT peptides) resuspended in RPMI+10%FBS (R10) with recombinant mouse IL-2 (rmIL-2) (R&D Systems) were added to each well. The final concentration of each peptide in the assay was 1µg/mL/peptide, and the final concentration of rmIL-2 was 0.1 ng/mL. R10 alone was used as a negative control and PMA+Ionomycin as a positive control. The ELISPOT plates were then placed into a humid 37°C with 5% CO2 incubator for 40-48 hours. After incubation, the plates were washed and IFN-γ capture antibody was added, followed by streptomycin horseradish peroxidase (strep-HRP). The plates were developed with commercially available 3-Amino-9-ethylcarbazole (AEC) substrate (Sigma-Aldrich). The observed spots were counted using an ELISPOT plate reader by ZellNet and the final data was reported as spot forming cells (SFC) per one million splenocytes.

### 2.11. Histopathology

At necropsy the left lung of hamsters was perfused with neutral-buffered formalin immediately after collection. Tissues were fixed in neutral-buffered formalin for a week, then placed into fresh neutral-buffered formalin before being transferred from containment level 3 to containment level 2 laboratory. Tissues were embeded, sectioned and stained with hematoxylin and eosin. Slides were examined by a board-certified pathologist.

### 2.12. Statistics

All statistical analyses were performed using GraphPad Prism 9 software (La Jolla, CA). Unless indicated, multiple comparison was done with Kruskall-Wallis test. The data were considered significant if p < 0.05. Geometric means with standard deviation are represented on graphs. No samples or animals were excluded from the analysis. Randomization was performed for the animal studies.

## 3. Results

### 3.1. Impact of SARS-CoV-2 S antigen design on expression and yield

Four constructs were designed based on the spike protein sequence of the SARS-CoV-2 Wuhan-Hu1 isolate and subcloned into expression plasmids for the production of eVLPs as described in Methods (Fig. 1a). To obtain a stabilized prefusion form of S (SP), the furin cleavage site of S, RRAR, was inhibited by mutation of the 3 arginines into a glycine and 2 serine (GSAS) and 2 proline substitutions were introduced at successive residues K986 and V987. Our previous work has demonstrated that the swap of the transmembrane cytoplasmic terminal domain (TMCTD) of CMV gB resulted in enhanced yields and immunogenicity of the gB glycoprotein presented on eVLPs [18]. Based on this data, two additional constructs, Native-VSVg (SG) and Stabilized Prefusion-VSVg (SPG) were designed by swapping the TMCTD of S with that of VSV-G.

**Fig. 1:**
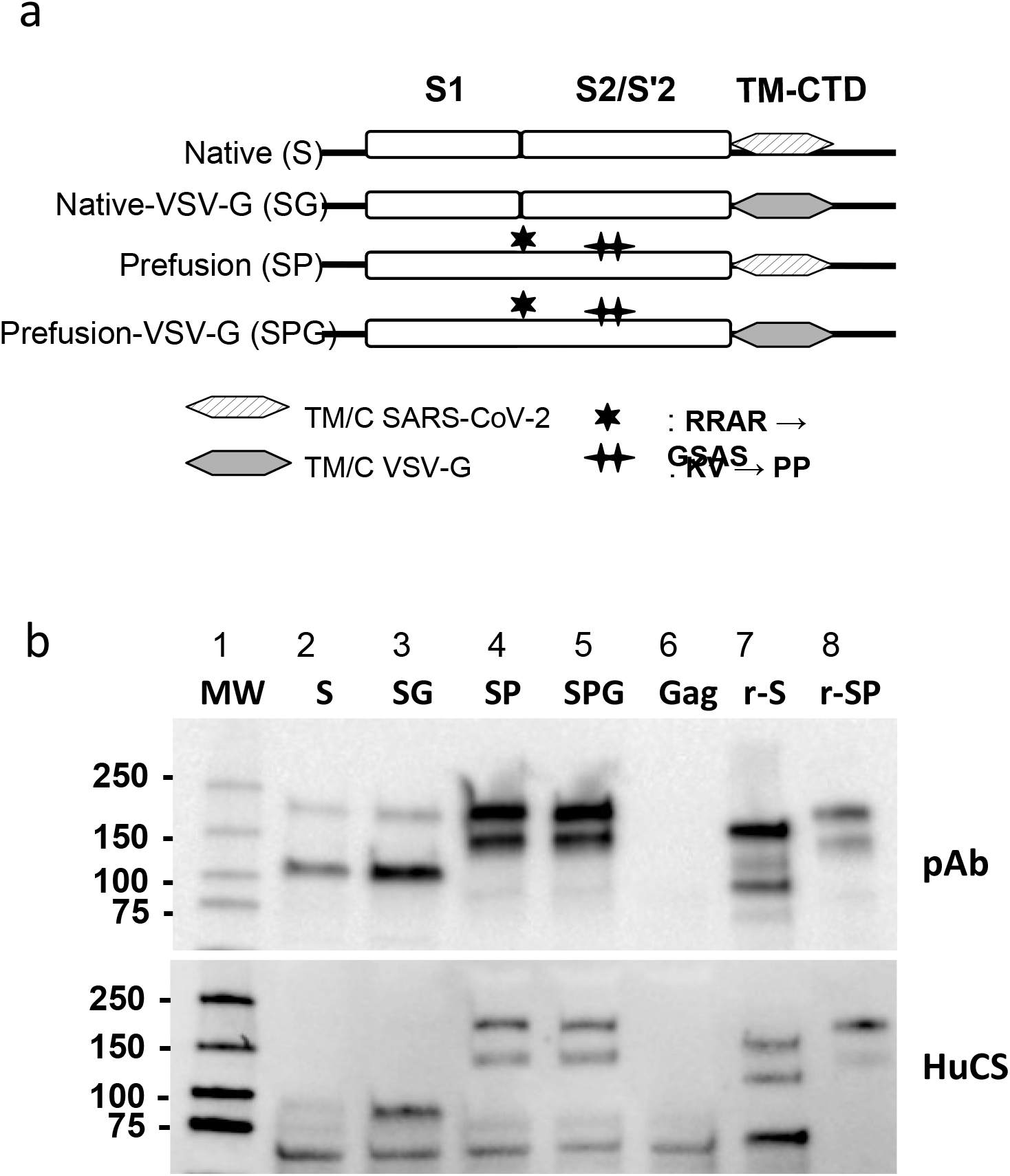
Constructs design and production of eVLPs expressing SARS-CoV-2 Spike protein. **(a)** Schematic representation of SARS-COV-2 S plasmid constructs. TM-CTD: Transmembrane cytoplasmic terminal domain. **(b)** Expression of SARS-CoV-2 S analyzed by Western-blot of SARS-CoV-2 eVLPs and recombinant proteins using a rabbit polyclonal Ab (pAb, upper panel) raised against SARS-CoV-2 RBD (Sinobiological) or COVID-19 convalescent human serum (HuCS, bottom panel). eVLPs produced with Gag plasmid only (Gag eVLPs) and recombinant SARS-CoV-2 S proteins were used as negative and positive controls respectively. r-S: recombinant native S, r-SP: recombinant prefusion S protein containing a mutated furin cleavage domain (RRAR → GSAS), replacement of 2 proline (KV → PP) and a trimerization domain.

Western blot analysis of eVLPs using a polyclonal Ab directed against the SARS-CoV-2 S receptor binding domain confirmed the processing of SARS-CoV-2 S during biosynthesis in HEK-293 cells as expected by the presence of the furin cleavage site in S1/S2 [6] (Fig. 1b, lane 2-3). Expression of S was slightly improved by the VSV-G swap in SG, and more dramatically enhanced by the inhibition of the cleavage sites in SP and SPG (Fig. 1b, lane 4-5). Overexpression of S in the prefusion forms showed a major band at 180 kDa, the size commonly described for uncleaved S180 kDa and an additional band around 150 kDa. The additional band around 150 kDa is reproducibly seen upon overexpression of uncleaved S, and most likely represents the S protein deprived of N-Glycosylation [21] that would occur because of overloading of the host cell machinery. Similar results were obtained after blotting with human convalescent sera.

Quantitative analysis of protein content in eVLP preparations showed that for a similar number of particles and comparable amounts of Gag protein, the amount of SARS-CoV-2 S protein was increased substantially with replacement of the TMCTD and by use of the stabilized prefusion construct, suggesting that the density of the S protein was enhanced using the VSV-G constructs (Table 1). The best yield was reproducibly obtained when producing the eVLPs expressing the prefusion-VSV-G form of S, with up to a 40-fold increase relative to eVLPs expressing native S.

**Table 1:** Optimisation of SARS-CoV-2 S protein yields by alteration of the sequence construct.

| Spike Construct | Gag<br>total amount<br>(mg) | SARS-CoV-2 S<br>total amount<br>(mg) | S to Gag ratio<br>(%) | Particle<br>number /mL |
| --- | --- | --- | --- | --- |
| Native (S) | 23 | 0.16 | 0.07 | $4.37 \times 10^{11}$ |
| Native-VSV-G (SG) | 19 | 0.5 | 0.26 | $4.37 \times 10^{11}$ |
| Prefusion (SP) | 32 | 0.23 | 0.71 | $4.37 \times 10^{11}$ |
| Prefusion-VSV-G (SPG) | 23 | 0.64 | 2.78 | $4.37 \times 10^{11}$ |

### 3.2. Impact of SARS-CoV-2 S antigen design on neutralizing antibody responses

Comparison to convalescent serum is commonly used as a benchmark to help evaluate immunogenicity and potential efficacy of Covid-19 candidate vaccines. However, a wide spectrum of Ab responses can be observed in recovering patients, ranging from barely detectable to very high levels, likely influenced by time since infection and severity of disease. To enable comparison across experiments, we obtained a cohort of 20 sera from COVID-19 confirmed convalescent patients with moderate COVID-19 symptoms who all recovered without specific treatment intervention or hospitalization. The cohort was separated into two groups of 10 samples according to high or low levels of Ab binding activity to recombinant SARS-CoV-2 S (Fig. 2a). Sera from each group were then pooled and tested for neutralizing activity (Fig. 2b). As expected, the pool of human sera showing higher levels of IgG titers against SARS-CoV-2 S had the highest neutralizing activity, which was consistent with previous observations [22]. To provide a robust benchmark with which to assess the immunogenicity of the vaccine candidates, only the high titer pooled sera was used to assess vaccine-induced responses in animals.

**Fig. 2:**
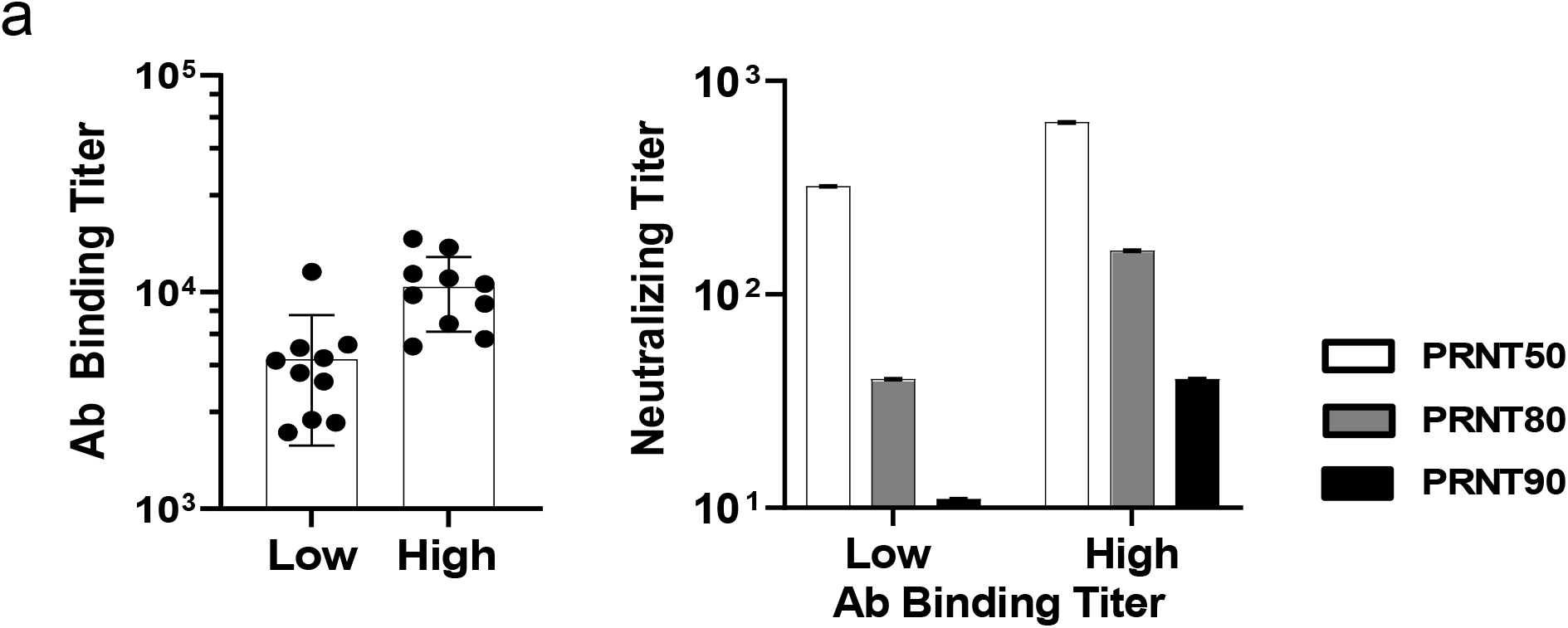
Measure of anti-SARS-CoV-2 Ab in COVID-19 convalescent human plasma. **(a)** Ab binding titers against SARS-CoV-2 S (S1+S2). Plasma samples were grouped according to initial screening by YHLO method in two groups Low and High Ab titers prior to be tested in the in-house ELISA against a recombinant S(S1+S2). **(b)** Neutralizing activity was measured by PRNT90 as described in Material and Methods. Results are represented as end point titers (EPT).

Humoral responses of the various types of SARS-CoV-2 eVLPs were evaluated in C57BL/6 mice that received 2 intraperitoneal injections at 3 week intervals (Fig. 3). The first injection of unmodified S presented on eVLPs induced levels of anti-SARS-CoV-2 S Ab binding titers similar to those in mice that received a recombinant trimerized prefusion S protein, but they were not associated with significant (90% or greater) neutralization activity as measured in a plaque reduction neutralization test (PRNT) (Fig. 3a-b). In contrast, a significant nAb response was induced by a single injection of eVLPs expressing prefusion SP or SPG, with PRNT90 end-point titers (EPTs) of 80 and 160 respectively. These values were higher than those observed with the human convalescent control pool (PRNT90 EPT of 50). All nAb responses were greatly enhanced by the second injection and reflected the responses that were observed prior to the boosting dose. Notably, all forms of SARS-CoV-2 S presented on eVLPs induced higher antibody titers than recombinant prefusion S protein, both in the levels of total IgG and neutralization activity, after one or two injections.

**Fig. 3:**
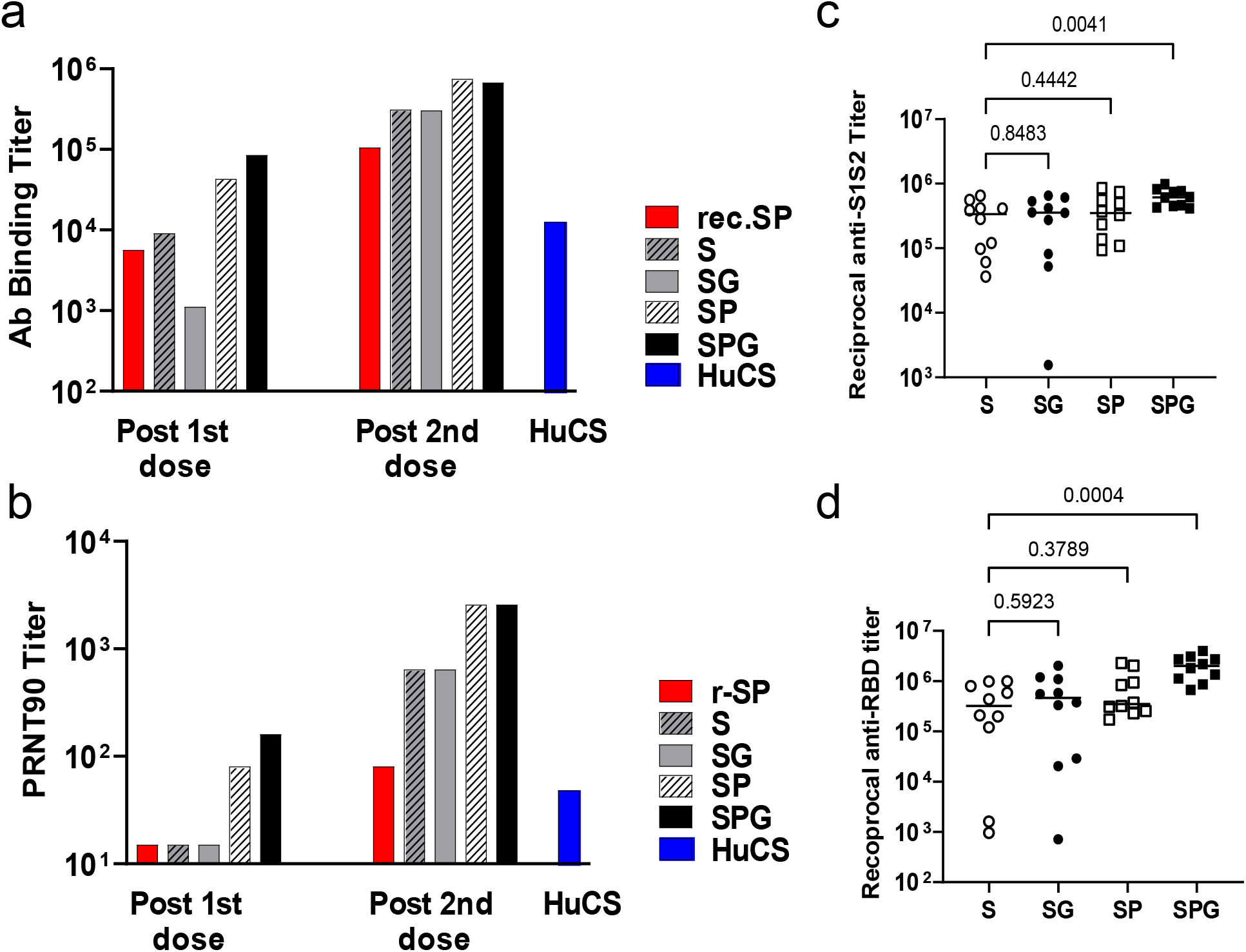
Immunogenicity of the various forms of SARS-CoV-2 S eVLPs in C57BL/6 mice. Four groups of 10 C57BL/6 mice received 2 injections of various forms of SARS-CoV-2 S eVLPs or recombinant SP at day 0 and 21 a indicated on legend, S: native S, SG: S with VSV-G tail, SP: prefusion S, SPG: prefusion S with VSV-G tail, r-SP: recombinant SP protein. Sera were collected 2 weeks after each injection. **(a)** Pooled sera from each group were analyzed for specific SARS-CoV-2 S(S1+S2) total IgG; results are represented as EPT corresponding to the first dilution that gave an OD 3-fold above background. **(b)** Pooled sera from each group were analyzed in PRNT assay with a 90% threshold (PRNT90) as described in Material and Methods. A pool of human sera from COVID-19 convalescent patients with moderate disease (HuCS) was used as reference. **(c-d)** Individual sera were analyzed in ELISA using recombinant SARS-CoV-2 S(S1+S2) protein (c) or recombinant SARS-CoV-2 RBD protein (d). P values from Kruskall-Wallis test comparing groups are indicated in c and d.

Individual mice sera obtained 14 days after the second injection of eVLPs were evaluated for the specificity of the Ab responses against the whole S1+S2 protein or the RBD (Fig. 3c-d). All immunized mice that received eVLPs showed robust anti-SARS-CoV-2 Ab responses either against a full length S1+S2 protein (Fig. 3c) or against the RBD protein (Fig. 3d). A more homogenous response was observed in mice that received the SPG eVLPs, with all Ab EPTs above 400,000 against S (5.6 Log 10), and above 650,000 against RBD (5.8 Log10).

### 3.3. Influence of adjuvants on antibody and T cell responses

A Th2-type response has been suggested to contribute to the “cytokine storm” associated with vaccine-induced severe lung pathologies [23,24]. In light of these results, we tested a variety of adjuvants that might enhance neutralizing Ab production while also promoting a balanced Th1/Th2 response. For this purpose, we compared formulation of eVLPs with Alum against a panel of adjuvants including MF59 and the adjuvant systems AS03 and AS04. We used SARS-CoV-2 native S eVLPs as they were less immunogenic than eVLPs expressing the prefusion form of the S protein and might better enable differences in the adjuvants to be observed. The various adjuvanted formulation of S eVLPs were compared to recombinant stabilized prefusion S protein (r-SP) formulated in Alum adjuvant, which was expected to induce a Th2-biased response [25]. Mice received two IP injections and Ab and T cell responses were measured 14 days after the second injection (Fig. 4). MF59 enhanced IFN-γ T cell responses compared to Alum (Fig. 4a) but induced similar Ab responses (Fig. 4b, c) and a comparable, balanced IgG2/IgG1 ratio (Fig. 4d). The AS03 and AS04 adjuvants also skewed responses towards a Th1-type T cell response. Most remarkably, while r-SP in Alum preferentially induced IgG1 Ab representative of a Th2 response, S-eVLPs induced balanced production of IgG1 and IgG2b indicating a balanced Th1/Th2 response (Fig. 4d).

**Fig. 4:**
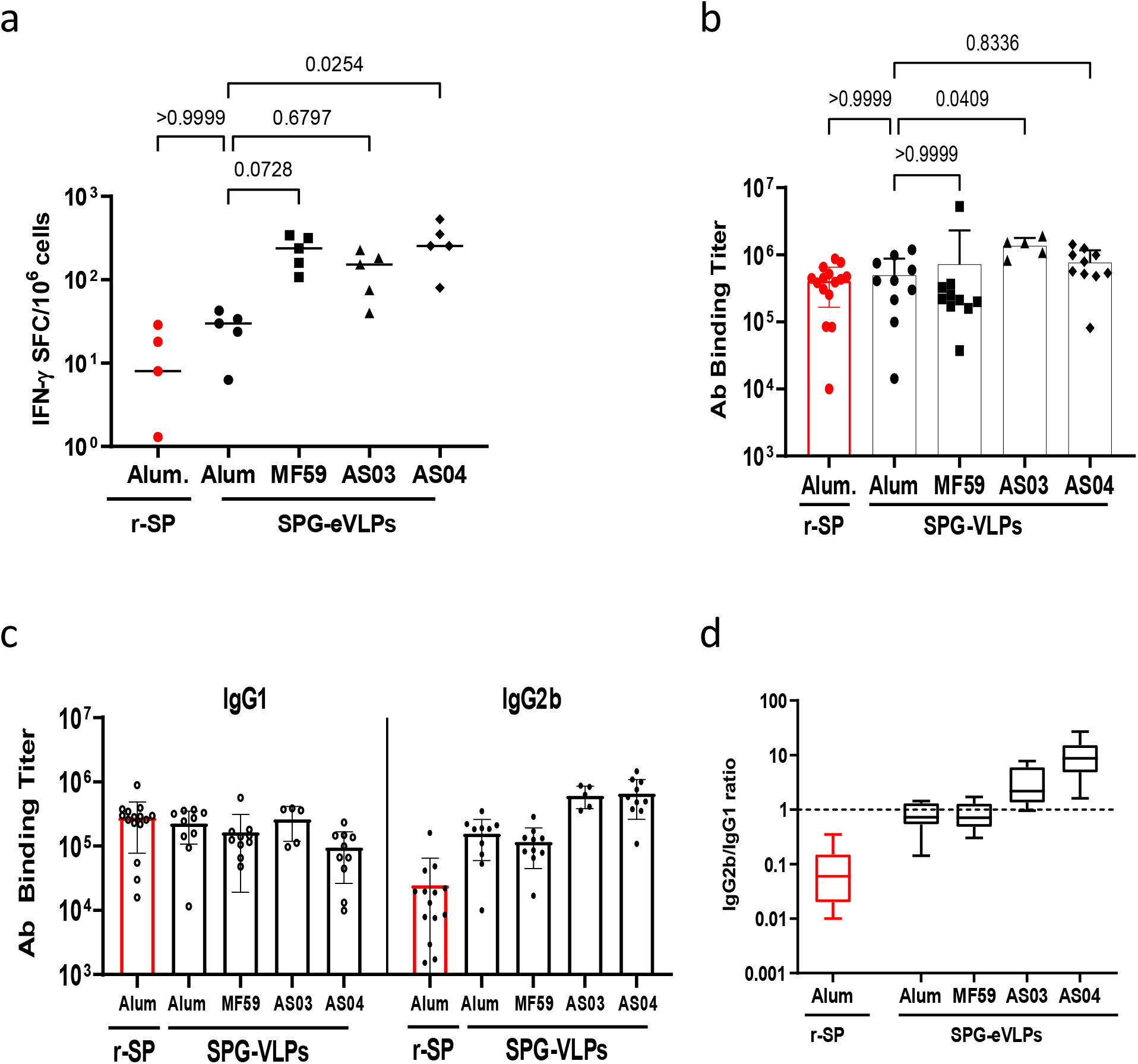
Influence of various adjuvant in the SARS-CoV-2 S eVLPs-mediated Ab and T cell response. At day 0 and 21, five groups of 10 C57BL/6 mice received 2 injections of S eVLPs in the presence of various adjuvants as indicated in legends and described in Material and Methods. Sera and splenocytes were collected 2 weeks after the second injection. **(a)** Numbers of IFNγ producing cells per million splenocytes collected 2 weeks after the second injection were measured by ELISpot using peptide pool covering the entire S(S1+S2) protein. **(b)** Total IgG were measured in ELISA against recombinant SARS-CoV-2 S(S1+S2) protein, results are represented as EPT. **(c-d)** Isotype usage was determined in individual sera by specific ELISA using HRP conjugate goat Ab against mouse IgG1 and IgG2. (c) Results are expressed as the ratio of IgG2b to IgG1. Results from Kruskall-Wallis comparison of groups are indicated.

### 3.4. Immunogenicity in mice of vaccine candidate VBI-2902a

Based on the results described above, we chose to evaluate the immunogenicity and potential efficacy of eVLPs expressing SPG protein formulated with Alum, named VBI-2902a, after one or two injections 21 days apart. Fourteen days after a single injection, sera from mouse immunized with VBI-2902a contained total anti-Spike IgG EPTs reaching geometric means of (4.8 Log 10) 54,891 that were associated with neutralizing PRNT90 titers of 365 (2.6 Log10). A second injection boosted Ab binding titers to 228,374 (5.4 Log10) with nAb titers of 1,079 (3.0 Log10) (Fig. 5a-b). Levels of nAb response were higher than those observed in sera from convalescent patients. Abs were preferentially directed against the RBD and S1 with only low binding to S2 (Fig. 5c). Mouse splenocytes collected 2 weeks after each immunization were stimulated *ex vivo* using two different peptide pools preferentially covering the S1 domain (pepmix 1) or the S2 domain (pepmix 2) respectively. Numbers of IFN-γ spot forming cells (Fig. 5d) suggested preferential T cell responses against the S1 domain of the spike protein rather than against the S2 domain. No major increases in T cell responses were observed after the second injection of VBI-2902a. Additionally, we observed that a single dose of VBI-2902a induced a sustained Ab response for at least 15 weeks without any drop in neutralization titers (Fig. 5e).

**Fig. 5:**
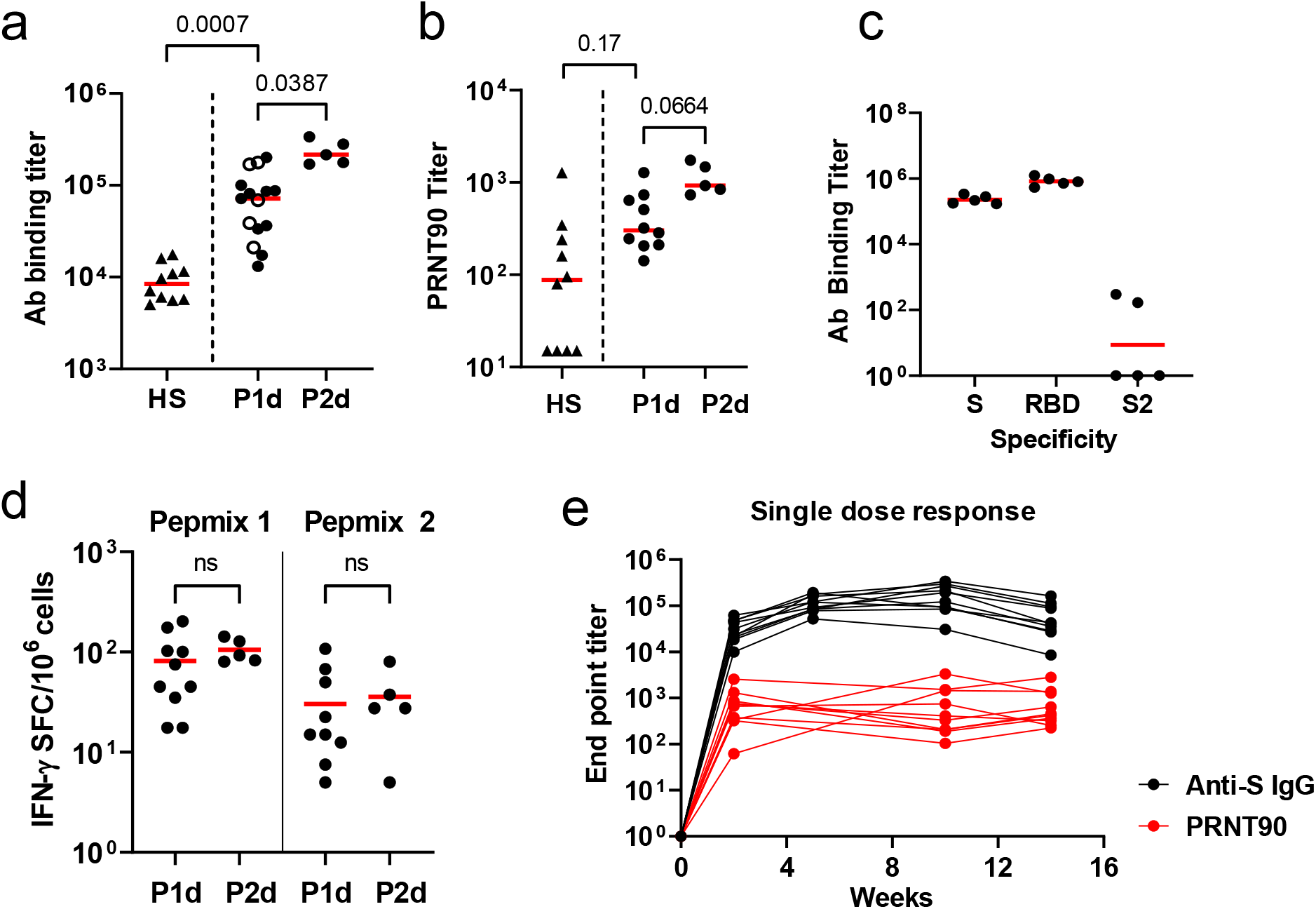
Immunogenicity of VBI-2902a and VBI-2902e in C57BL/6 mice. **(a-d)** Two groups of 10 mice were immunized twice at 3 weeks interval with VBI-2902a or VBI-2902e containing 0.2 µg of S protein. Blood was collected 2 weeks after each injection, P1d: post 1^st^ dose, P2d: post 2^nd^ dose. **(a)** Ab binding titer against recombinant S(S1+S2) compared to human convalescent sera, measured by ELISA, **(b)** neutralization end point titers measured by PRNT90, **(c)** Ab binding titers against recombinant S(S1+S2), recombinant RBD or recombinant S2 measured by ELISA in sera after the 2^nd^ dose. Results from Kruskall-Wallis comparison of groups are indicated for A and B. **(d)** Numbers of IFNγ producing cells per million splenocytes collected 2 weeks after each injection were measured by ELISpot using Pepmix 1 or Pepmix 2 preferentially covering SARS-CoV-2 S1 domain or tS2 domain respectively. **(e)** Kinetic of the humoral response after single injection of VBI-2902a calculated as end point titer determined in ELISA and PRNT90.

### 3.5. Protective efficacy of VBI-2902a in Syrian Golden Hamsters

The protective efficacy of VBI-2902a was examined in Syrian Gold hamsters. SARS-CoV-2 infection in Syrian Gold hamsters resembles features found in humans with moderate COVID-19 and is characterized by a rapid weight loss starting 2 days post infection (dpi) [26,27]. Two immunization regimens were compared. Regimen II consisted of two IM injections of VBI-2902a or saline at 3 weeks interval whereas Regimen I consisted of a single dose injection of VBI-2902a or saline (Fig. 6a). Three weeks after the last injection (day 42 in Regimen II and day 21 in Regimen I), all animals were inoculated intranasally with 1×10^5^ TCID50 of SARS-CoV-2 per animal and monitored daily for weight change, general health and behavior.

After a single injection of VBI-2902a the levels of anti-S IgG rapidly increased in the serum of immunized animals with EPTs reaching 1-2×10^3^ (Fig. 6b). The second injection enhanced these levels approximately 10-fold to reach EPTs of 2-3×10^4^ at day 35, which translated into robust neutralization titers of over 10^3^ EPT as measured by PRNT90 (Fig. 6c). In the single dose regimen, the neutralization activity (GeoMean 69), was increased 250-fold to 1725 within 3 days after exposure to the virus.

Animals in all groups lost 2-4% of body weight 2 days post infection (2dpi). Animals in the saline control groups continued to lost weight until an average 15% loss at 7dpi, before gradually regaining weight (Fig. 7a-b). In marked contrast, none of the hamsters immunized with two doses of VBI-2902a lost any further weight after 2dpi, regaining normal weight by 7dpi, demonstrating robust protection against SARS-CoV-2 disease. In the single dose regimen, the majority of the animals regained body weight after 3dpi instead of 2dpi, suggesting slightly delayed but significant protection against disease.

**Fig. 6:**
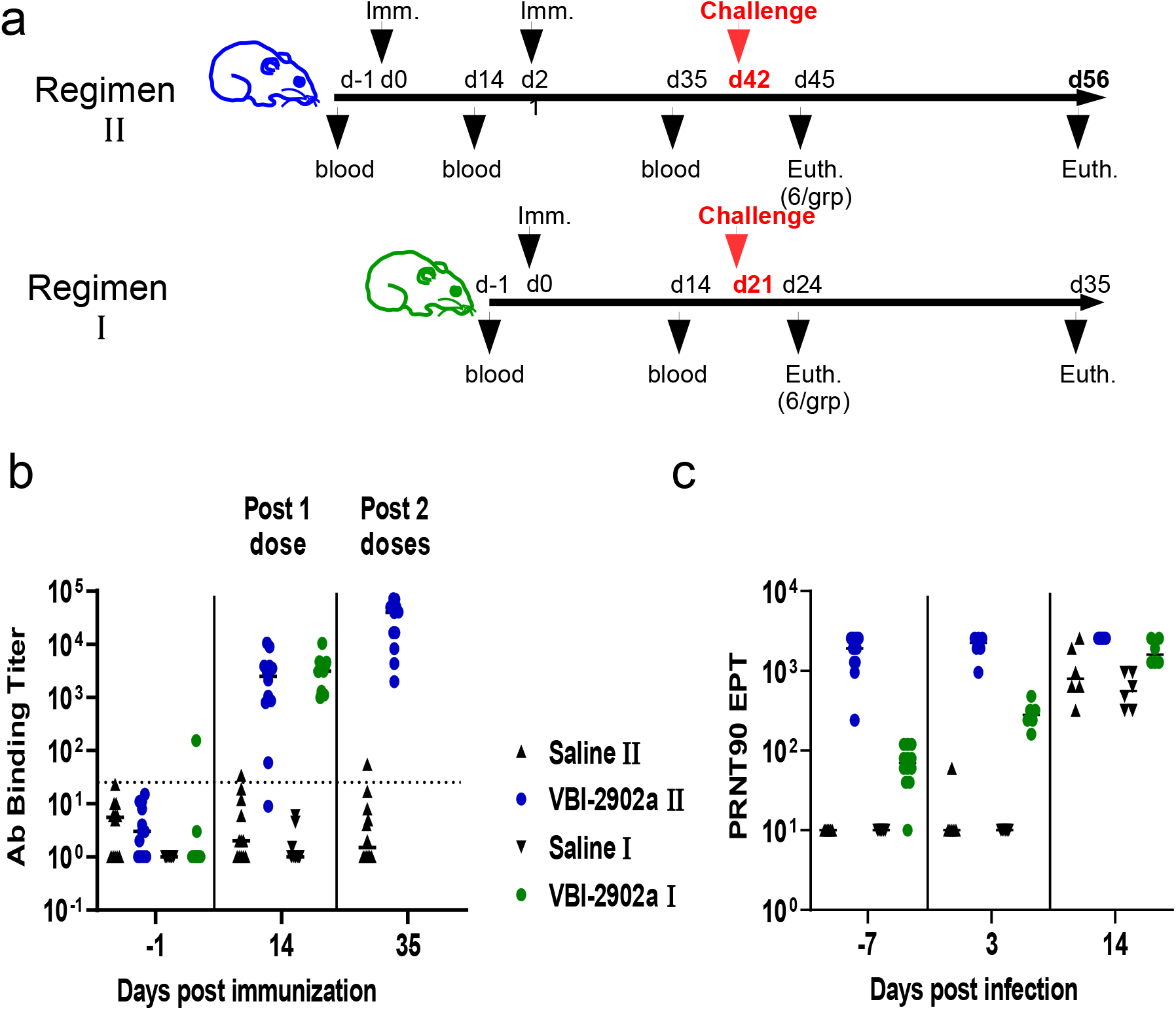
Immunogenicity of VBI-2902a or VBI-2902e in Syrian gloden hamsters. **(a)** Schematic representation of the challenge experiments. Each challenge experiment used 2 groups of 15 Syrian gold hamsters. In regimen II, animals received 2 IM injections of VBI-2902a (2µg of S per dose) or placebo saline buffer administered at 3 weeks interval. In regimen I, animals received a single injection of VBI-2902a or Saline buffer. Blood was collected 2 weeks after each injection. Three weeks after the last injection corresponding to day 42 in regimen II and day 21 in regimen I, hamsters were exposed to SARS-CoV-2 at 1×10^5^ TCID50 per animal via both nares. At 3 days post infection (dpi), 6 animals per groups were sacrificed for viral load analysis. The remaining animals were clinically evaluated daily until end of study at 14dpi. **(b)** Anti-SARS-CoV-2 S(S1+S2) total IgG EPT measured by ELISA 2 weeks after each immunization. **(c)** Neutralization activity was measured by PRNT90 in immunized groups; results are represented as PRNT90 EPT.

**Fig. 7:**
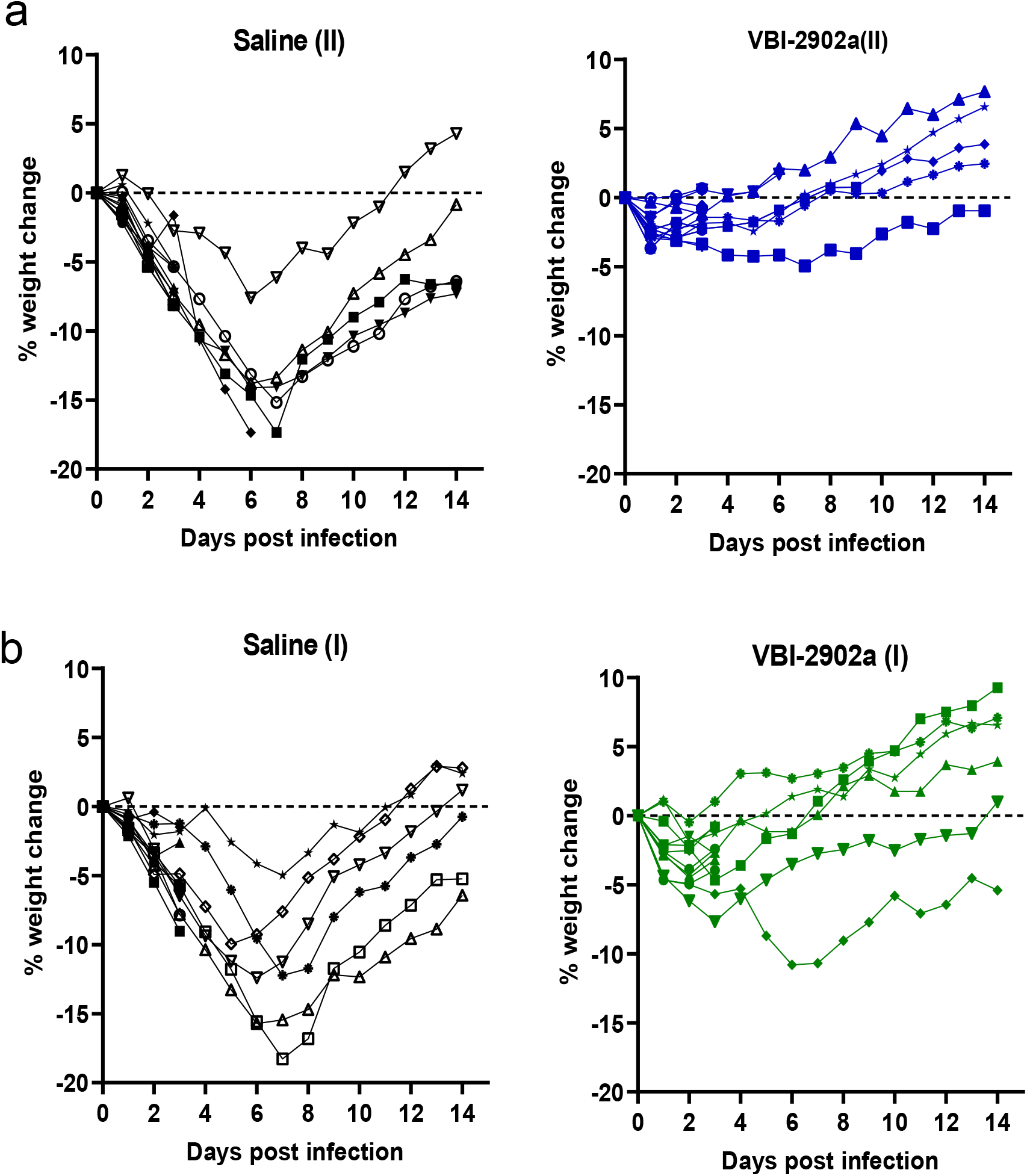
Weight change ofhamsters after exposure to SARS-CoV-2. Hamsters from experiment described in fig. 6 were monitored daily for weight change. Results are represented for each animal in each groups as kinetic of weight change from SARS-CoV-2 exposure to day 9 after infection. **(a)** represents the weight change observed in the 2-dose regimen (II), **(b)** represent the weight change observed in the single dose regimen (I).

At 3dpi, hamsters vaccinated with either one or two doses of VBI-2902a had greatly decreased viral RNA copy numbers in lungs (Fig. 8a). Two doses of VBI-2902a resulted in a 5 Log decrease in viral load in the cranial lobe and a 4 Log decrease in the caudal lobe relative to non-immunized animals; a single dose of vaccine induced a 2 Log decrease in the cranial lobe and a 4 Log decrease in the caudal lobe. The viral load values observed in lungs were inversely correlated with the neutralization measured as PRNT90 (Fig. c-<u>d</u>). More viral RNA was found in nasal turbinates, which may have included residual viral inoculum as suggested previously [27,28]. Data from prior studies also suggested an extended persistence of the virus in nasal turbinates while bearly detectable in the lung [26]. Both vaccine regimens protected against the development of lung pathology as indicated by reductions of the lung to body weight ratio (Fig. 9a-b) and histological analysis of the lungs (Fig. 9c-d).

**Fig. 8:**
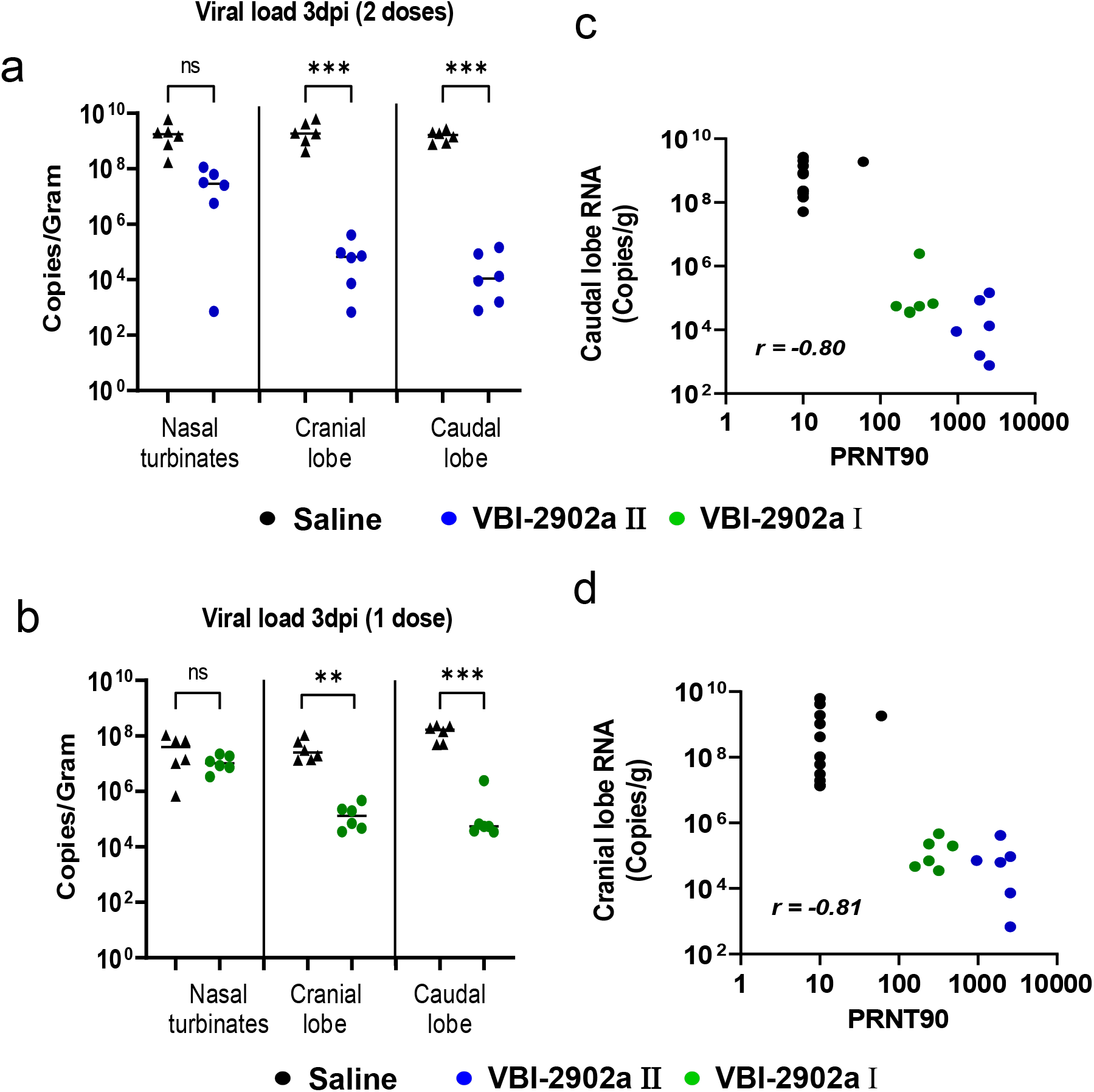
Viral load analysis in SARS-CoV-2 infected hamsters. **(a-b)** At 3dpi, qRT-PCR assays were performed on RNA from samples of nasal washes, lung tissues (cranial and caudal lobes) using SARS-CoV-2 specific primers. Results were expressed as copy number per gram of tissue sample. **(c-d)** Correlation analysis of viral loads measured in lung caudal lobe and PRNT90.

**Fig. 9:**
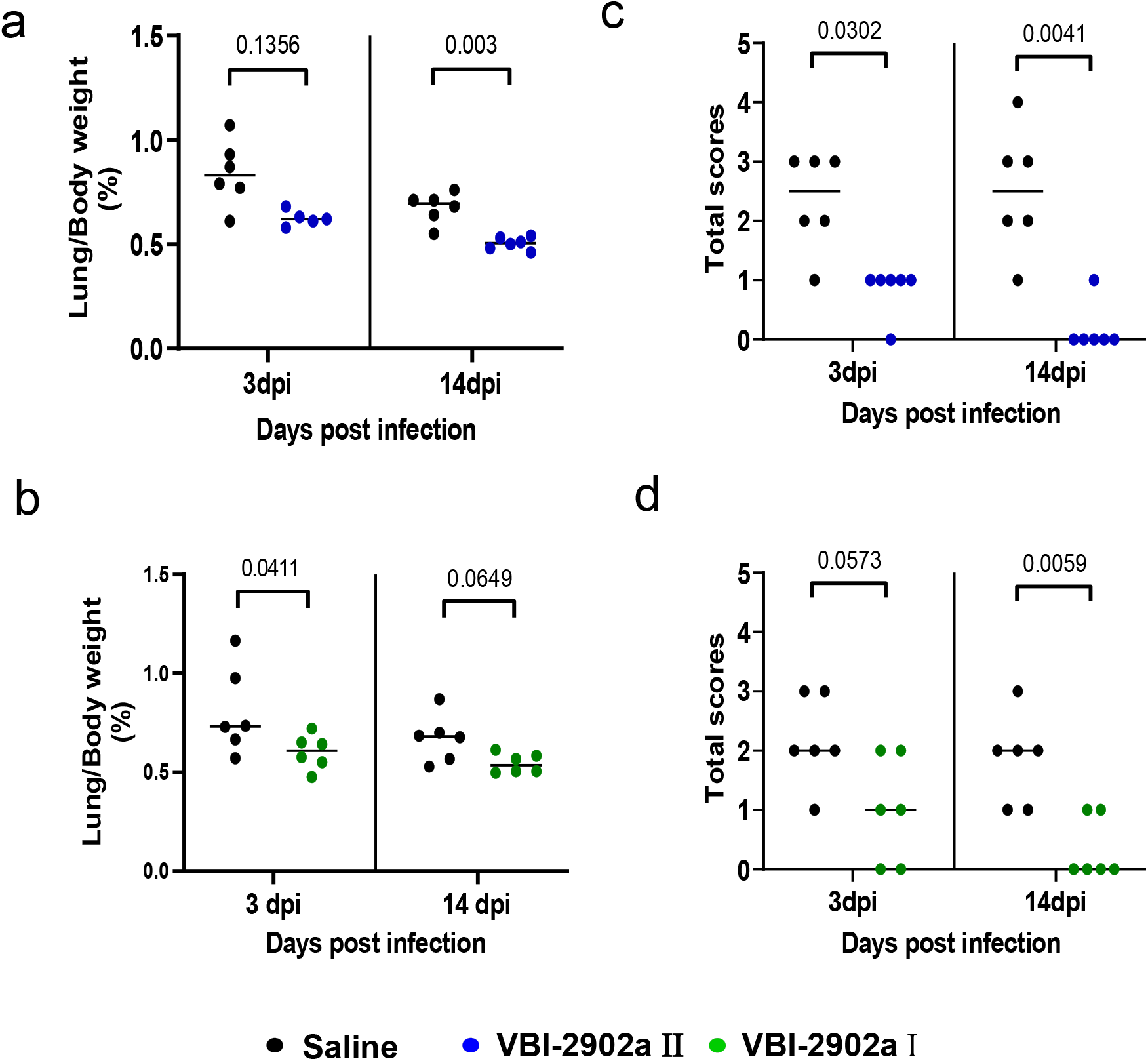
Clinical evaluation of lung pathology in immunized hamsters challenged by SARS-CoV-2 virus. **(a-b)** lung to body weight ratio in hamsters at 3dpi and 14dpi. **(c-d)** Histopathology severity analysis of hamster lungs at 3dp and 14 dpi. Scores were evaluated on a scale from 0 to 4 as follow: 0, no microscopic lesions; 1, slight or questionable pneumonia; 2, clearly present, but not conspicuously so; 3, moderate pneumonia; 4, severe pneumonia. Statistics were performed using Kruskal-Wallis non-parametric test followed by Dunn’ multiple comparisons test. Adjusted p values are shown.

## 4. Discussion

The unprecedented urgency for a safe COVID-19 vaccine that can confer protection as quickly as possible with as few doses as possible is evident as regulatory agencies and vaccine manufactures have discussed the risks and benefits of delaying planned second doses of currently available COVID-19 vaccines to enable immunization of a greater number of individuals as quickly as possible [29,30]. We have previously demonstrated that expression of proteins on the surface of eVLPs dramatically enriches for neutralizing antibody, the presumed correlate of protection against SARS-CoV-2, relative to recombinant proteins [18]. Accordingly, we evaluated both conformations of the SARS-CoV-2 S protein as well as a variety of adjuvants in an effort to identify a COVID-19 vaccine candidate with the potential to confer protection after a single dose.

The eVLPs particles were pseudotyped with SARS-CoV-2 unmodified S protein but expressed low amounts of S that were not suitable for upscaled production. We therefore designed a modified prefusion form of S that resulted in both dramatic increases in yields and enhancement of the nAb response compared to native S. SPG eVLPs induced high titers of RBD Ab binding titers associated with robust neutralizing responses in mice at levels that were much higher than those observed with a recombinant prefusion S protein. Indeed, 14 days after a single dose of SPG eVLPs formulated with aluminum phosphate nAb titers exceeded those associated with high titer COVID-19 covalescent sera, persisted and were undimished for at least 3 months. The potency observed after a single dose of VBI-2902a appears superior to what has been observed after 2 doses in the same strain of mice with an mRNA vaccine that has received Emergency Use Authorization [12], further demonstrating the strong potency of this vaccine candidate. In a hamster challenge model VBI-2902a demonstrated robust efficacy against clinical disease and lung inflammation. While two doses showed greater efficacy, a single dose clearly conferred protective benefit.

The value of eVLP expression of the modified SP protein is consistent with prior reports which demonstrated that an anchored version of a stabilized prefusion S antigen provided optimal induction of protective nAbs in Rhesus macaques [13]. Our construct differed from the previously described S-2P [13, 31] by using the VSV-G transmembrane cytoplasmic domain to replace that of S, instead of a C-terminal T4 fibritin trimerization domain. Based on previous experience and published data [18,19], we hypothesized that the use of VSV-G tail and expression in the phospholipid membrane of eVLPs would result in natural trimerization of the spike ectodomains providing optimal presentation of neutralization epitopes. The use of the VSV-G tail has been shown to enhance expression and localization of viral glycoproteins at the phospholipid envelop of the particles [32,33].

Aluminum salt adjuvants have a long history of safety and are a component of approved VLP-based vaccines such as Gardasil® against HPV [34] and Engerix B® against HBV [35]. Nevertheless, theoretical concerns have been raised about the use of an aluminum-based adjuvant with a SARS-CoV-2 vaccine and the potential for Th2-mediated enhanced lung pathology [36,37]. Subsequent studies have demonstrated that non-neutralizing antibodies against structural proteins were responsible for the pathology observed in preclinical models [38]. Use of eVLP presentation of an optimized form of the SARS-CoV-2 S protein resulted in a highly potent and focused neutralizing antibody response which avoided any evidence of disease enhancement or increased lung inflammation. In a hamster challenge model VBI-2902a demonstrated efficacy and ability to suppress lung inflammation. While two doses showed better potency, the single dose also conferred protective benefit indicated by comparable results in terms of lung inflammation.

Moreover, compared to a clear Th2-biased profile observed in response to recombinant prefusion stabilized S protein in Alum, the similar prefusion S construct induced a balanced Th1/Th2 response when presented by eVLPs (Fig. 4d). The balanced production of IgG2/IgG1 antibody isotypes after VBI-2902a immunization was comparable with those described in response to the recently emergency use authorized vaccine Ad26.COV2.S [25]. These results emphasize an important difference in the quality of the antibody response when immunizing with soluble, recombinant versus particulate forms of vaccine antigens.

The VBI-2902a vaccine candidate addresses several issues that have thus far hindered the speed and extent of vaccination with currently available COVID-19 vaccines. This includes the need to administer multiple doses and the need for storage, transport, and distribution of the vaccine at freezing temperatures not typically required for prophylactic vaccines. VBI-2902a received approval from Health Canada to initiate its ongoing Phase I/II clinical study (NCT04773665) to assess its potential for one dose immunogenicity and potential efficacy.

## Acknowledgment

The authors want to thank Adam Asselin, Matthew Yorke, Teresa Daoud, Lanjian (Isabel) Yang, Rebecca Wang, Gillian Lampkin (VBI vaccines) for outstanding technical support;Traian Sulea (NRC) for discussions on construct design, NRC Animal Resources Group and the VIDO Saskatchewan team for remarkable care with animal experiments as well as Ammon Ding and Echo Wu (Genescript) for their dedication in plasmid preparation. All the people cited above contributed to the success of the study by the excellence of their work.

## Abbreviations

eVLP,: enveloped virus-like particules;
CoV,: coronavirus;
RBD,: receptor binding domain;
TMCTD,: transmembrane cytoplasmic terminal domain;
Ab,: antibody;
nAb,: neutralizing antibody;
MLV,: murine leukemia virus;
ELISA,: enzyme-linked-immuno-sorbent-assay;
PRNT,: plaque reduction neutralization test;
EPT,: end-point titer; Alum, aluminum;
ELISPOT,: Enzyme Linked ImmunoSpot Test;
IP,: IntraPeritoneal;
IM,: IntraMuscular;
NRC,: National Research Council Canada;
VIDO,: Vaccine and Infectious Disease Organization

## References

[1] WHO weekly reports, https://www.who.int/emergencies/diseases/novel-coronavirus-2019/situation-reports

[2] To KK, Hung IF, Ip JD, Chu AW, Chan WM, Tam AR, Fong CH, Yuan S, Tsoi HW, Ng AC, Lee LL, Wan P, Tso E, To WK, Tsang D, Chan KH, Huang JD, Kok KH, Cheng VC, Yuen KY. COVID-19 re-infection by a phylogenetically distinct SARS-coronavirus-2 strain confirmed by whole genome sequencing. Clin Infect Dis. 2020 Aug 25:ciaa1275. doi: 10.1093/cid/ciaa1275. Epub ahead of print. PMID: 32840608; PMCID: PMC7499500.

[3] Tillett RL, Sevinsky JR, Hartley PD, Kerwin H, Crawford N, Gorzalski A, Laverdure C, Verma SC, Rossetto CC, Jackson D, Farrell MJ, Van Hooser S, Pandori M. Genomic evidence for reinfection with SARS-CoV-2: a case study. Lancet Infect Dis. 2021 Jan;21(1):52–58. doi: 10.1016/S1473-3099(20)30764-7. Epub 2020 Oct 12. PMID: 33058797; PMCID: PMC7550103.

[4] Lavine JS, Bjornstad ON, Antia R. Immunological characteristics govern the transition of COVID-19 to endemicity. Science. 2021 Jan 12:eabe6522. doi: 10.1126/science.abe6522. Epub ahead of print. PMID: 33436525.

[5] Li F. Structure, Function, and Evolution of Coronavirus Spike Proteins. Annu Rev Virol. 2016 Sep 29;3(1):237–261. doi: 10.1146/annurev-virology-110615-042301. Epub 2016 Aug 25. PMID: 27578435; PMCID: PMC5457962.

[6] Walls AC, Park YJ, Tortorici MA, Wall A, McGuire AT, Veesler D. Structure, Function, and Antigenicity of the SARS-CoV-2 Spike Glycoprotein. Cell. 2020 Apr 16;181(2):281-292.e6. doi: 10.1016/j.cell.2020.02.058. Epub 2020 Mar 9. Erratum in: Cell. 2020 Dec 10;183(6):1735. PMID: 32155444; PMCID: PMC7102599.

[7] Walls AC, Tortorici MA, Snijder J, Xiong X, Bosch BJ, Rey FA, Veesler D. Tectonic conformational changes of a coronavirus spike glycoprotein promote membrane fusion. Proc Natl Acad Sci U S A. 2017 Oct 17;114(42):11157–11162. doi: 10.1073/pnas.1708727114. Epub 2017 Oct 3. PMID: 29073020; PMCID: PMC5651768.

[8] Coutard B, Valle C, de Lamballerie X, Canard B, Seidah NG, Decroly E. The spike glycoprotein of the new coronavirus 2019-nCoV contains a furin-like cleavage site absent in CoV of the same clade. Antiviral Res. 2020 Apr;176:104742. doi: 10.1016/j.antiviral.2020.104742. Epub 2020 Feb 10. PMID: 32057769; PMCID: PMC7114094.

[9] McLellan JS, Chen M, Leung S, Graepel KW, D. X, Yang Y, Zhou T, Baxa U, Yasuda E, Beaumont T, Kumar A, Modjarrad K, Zheng Z, Zhao M, Xia N, Kwong PD, Graham BS. Structure of RSV fusion glycoprotein trimer bound to a prefusion-specific neutralizing antibody. Science. 2013 May 31;340(6136):1113–7. doi: 10.1126/science.1234914. Epub 2013 Apr 25. PMID: 23618766; PMCID: PMC4459498. Pallesen J, Wang N, Corbett KS, Wrapp D, Kirchdoerfer RN, Turner HL, Cottrell CA, Becker MM, Wang L, Shi W, Kong WP, Andres EL, Kettenbach AN, Denison MR, Chappell JD, Graham BS, Ward AB, McLellan JS. Immunogenicity and structures of a rationally designed prefusion MERS-CoV spike antigen. Proc Natl Acad Sci U S A. 2017 Aug 29;114(35):E7348–E7357. doi: 10.1073/pnas.1707304114. Epub 2017 Aug 14. PMID: 28807998; PMCID: PMC5584442.

[10] Wrapp D, Wang N, Corbett KS, Goldsmith JA, Hsieh CL, Abiona O, Graham BS, McLellan JS. Cryo-EM structure of the 2019-nCoV spike in the prefusion conformation. Science. 2020 Mar 13;367(6483):1260–1263. doi: 10.1126/science.abb2507. Epub 2020 Feb 19. PMID: 32075877; PMCID: PMC7164637.

[11] Corbett KS, Edwards DK, Leist SR, Abiona OM, Boyoglu-Barnum S, Gillespie RA, Himansu S, Schäfer A, Ziwawo CT, DiPiazza AT, Dinnon KH, Elbashir SM, Shaw CA, Woods A, Fritch EJ, Martinez DR, Bock KW, Minai M, Nagata BM, Hutchinson GB, Wu K, Henry C, Bahl K, Garcia-Dominguez D, Ma L, Renzi I, Kong WP, Schmidt SD, Wang L, Zhang Y, Phung E, Chang LA, Loomis RJ, Altaras NE, Narayanan E, Metkar M, Presnyak V, Liu C, Louder MK, Shi W, Leung K, Yang ES, West A, Gully KL, Stevens LJ, Wang N, Wrapp D, Doria-Rose NA, Stewart-Jones G, Bennett H, Alvarado GS, Nason MC, Ruckwardt TJ, McLellan JS, Denison MR, Chappell JD, Moore IN, Morabito KM, Mascola JR, Baric RS, Carfi A, Graham BS. SARS-CoV-2 mRNA vaccine design enabled by prototype pathogen preparedness. Nature. 2020 Oct;586(7830):567–571. doi: 10.1038/s41586-020-2622-0. Epub 2020 Aug 5. PMID: 32756549; PMCID: PMC7581537.

[12] Mercado NB, Zahn R, Wegmann F, Loos C, Chandrashekar A, Yu J, Liu J, Peter L, McMahan K, Tostanoski LH, He X, Martinez DR, Rutten L, Bos R, van Manen D, Vellinga J, Custers J, Langedijk JP, Kwaks T, Bakkers MJG, Zuijdgeest D, Rosendahl Huber SK, Atyeo C, Fischinger S, Burke JS, Feldman J, Hauser BM, Caradonna TM, Bondzie EA, Dagotto G, Gebre MS, Hoffman E, Jacob-Dolan C, Kirilova M, Li Z, Lin Z, Mahrokhian SH, Maxfield LF, Nampanya F, Nityanandam R, Nkolola JP, Patel S, Ventura JD, Verrington K, Wan H, Pessaint L, Van Ry A, Blade K, Strasbaugh A, Cabus M, Brown R, Cook A, Zouantchangadou S, Teow E, Andersen H, Lewis MG, Cai Y, Chen B, Schmidt AG, Reeves RK, Baric RS, Lauffenburger DA, Alter G, Stoffels P, Mammen M, Van Hoof J, Schuitemaker H, Barouch DH. Single-shot Ad26 vaccine protects against SARS-CoV-2 in rhesus macaques. Nature. 2020 Oct;586(7830):583–588. doi: 10.1038/s41586-020-2607-z. Epub 2020 Jul 30. PMID: 32731257; PMCID: PMC7581548.

[13] Anderson EJ, Rouphael NG, Widge AT, Jackson LA, Roberts PC, Makhene M, Chappell JD, Denison MR, Stevens LJ, Pruijssers AJ, McDermott AB, Flach B, Lin BC, Doria-Rose NA, O’Dell S, Schmidt SD, Corbett KS, Swanson PA 2nd, Padilla M, Neuzil KM, Bennett H, Leav B, Makowski M, Albert J, Cross K, Edara VV, Floyd K, Suthar MS, Martinez DR, Baric R, Buchanan W, Luke CJ, Phadke VK, Rostad CA, Ledgerwood JE, Graham BS, Beigel JH; mRNA-1273 Study Group. Safety and Immunogenicity of SARS-CoV-2 mRNA-1273 Vaccine in Older Adults. N Engl J Med. 2020 Dec 17;383(25):2427–2438. doi: 10.1056/NEJMoa2028436. Epub 2020 Sep 29. PMID: 32991794; PMCID: PMC7556339.

[14] Walsh EE, Frenck RW Jr, Falsey AR, Kitchin N, Absalon J, Gurtman A, Lockhart S, Neuzil K, Mulligan MJ, Bailey R, Swanson KA, Li P, Koury K, Kalina W, Cooper D, Fontes-Garfias C, Shi PY, Türeci Ö, Tompkins KR, Lyke KE, Raabe V, Dormitzer PR, Jansen KU, Şahin U, Gruber WC. Safety and Immunogenicity of Two RNA-Based Covid-19 Vaccine Candidates. N Engl J Med. 2020 Dec 17;383(25):2439–2450. doi: 10.1056/NEJMoa2027906. Epub 2020 Oct 14. PMID: 33053279; PMCID: PMC7583697.

[15] Roldão A, Mellado MC, Castilho LR, Carrondo MJ, Alves PM. Virus-like particles in vaccine development. Expert Rev Vaccines. 2010 Oct;9(10):1149–76. doi: 10.1586/erv.10.115. PMID: 20923267.

[16] Bachmann MF, Rohrer UH, Kündig TM, Bürki K, Hengartner H, Zinkernagel RM. The influence of antigen organization on B cell responsiveness. Science. 1993 Nov 26;262(5138):1448–51. doi: 10.1126/science.8248784. PMID: 8248784.

[17] Kirchmeier M, Fluckiger AC, Soare C, Bozic J, Ontsouka B, Ahmed T, Diress A, Pereira L, Schödel F, Plotkin S, Dalba C, Klatzmann D, Anderson DE. Enveloped virus-like particle expression of human cytomegalovirus glycoprotein B antigen induces antibodies with potent and broad neutralizing activity. Clin Vaccine Immunol. 2014 Feb;21(2):174–80. doi: 10.1128/CVI.00662-13. Epub 2013 Dec 11. PMID: 24334684; PMCID: PMC3910943.

[18] Garrone P, Fluckiger AC, Mangeot PE, Gauthier E, Dupeyrot-Lacas P, Mancip J, Cangialosi A, Du Chéné I, LeGrand R, Mangeot I, Lavillette D, Bellier B, Cosset FL, Tangy F, Klatzmann D, Dalba C. A prime-boost strategy using virus-like particles pseudotyped for HCV proteins triggers broadly neutralizing antibodies in macaques. Sci Transl Med. 2011 Aug 3;3(94):94ra71. doi: 10.1126/scitranslmed.3002330. PMID: 21813755.

[19] Sun Z, Ren K, Zhang X, Chen J, Jiang Z, Jiang J, Ji F, Ouyang X, Li L. Mass Spectrometry Analysis of Newly Emerging Coronavirus HCoV-19 Spike Protein and Human ACE2 Reveals Camouflaging Glycans and Unique Post-Translational Modifications. Engineering (Beijing). 2020 Aug 30. doi: 10.1016/j.eng.2020.07.014. Epub ahead of print. PMID: 32904601; PMCID: PMC7456593.

[20] Ni L, Ye F, Cheng ML, Feng Y, Deng YQ, Zhao H, Wei P, Ge J, Gou M, Li X, Sun L, Cao T, Wang P, Zhou C, Zhang R, Liang P, Guo H, Wang X, Qin CF, Chen F, Dong C. Detection of SARS-CoV-2-Specific Humoral and Cellular Immunity in COVID-19 Convalescent Individuals. Immunity. 2020 Jun 16;52(6):971-977.e3. doi: 10.1016/j.immuni.2020.04.023. Epub 2020 May 3. PMID: 32413330; PMCID: PMC7196424.

[21] Peeples L. News Feature: Avoiding pitfalls in the pursuit of a COVID-19 vaccine. Proc Natl Acad Sci U S A. 2020 Apr 14;117(15):8218–8221. doi: 10.1073/pnas.2005456117. Epub 2020 Mar 30. PMID: 32229574; PMCID: PMC7165470.

[22] Roncati L, Nasillo V, Lusenti B, Riva G. Signals of T h 2 immune response from COVID-19 patients requiring intensive care. Ann Hematol. 2020 Jun;99(6):1419–1420. doi: 10.1007/s00277-020-04066-7. Epub 2020 May 8. PMID: 32382776; PMCID: PMC7205481.

[23] Bos R, Rutten L, van der Lubbe JEM, Bakkers MJG, Hardenberg G, Wegmann F, Zuijdgeest D, de Wilde AH, Koornneef A, Verwilligen A, van Manen D, Kwaks T, Vogels R, Dalebout TJ, Myeni SK, Kikkert M, Snijder EJ, Li Z, Barouch DH, Vellinga J, Langedijk JPM, Zahn RC, Custers J, Schuitemaker H. Ad26 vector-based COVID-19 vaccine encoding a prefusion-stabilized SARS-CoV-2 Spike immunogen induces potent humoral and cellular immune responses. NPJ Vaccines. 2020 Sep 28;5:91. doi: 10.1038/s41541-020-00243-x. PMID: 33083026; PMCID: PMC7522255.

[24] Chan JF, Zhang AJ, Yuan S, Poon VK, Chan CC, Lee AC, Chan WM, Fan Z, Tsoi HW, Wen L, Liang R, Cao J, Chen Y, Tang K, Luo C, Cai JP, Kok KH, Chu H, Chan KH, Sridhar S, Chen Z, Chen H, To KK, Yuen KY. Simulation of the Clinical and Pathological Manifestations of Coronavirus Disease 2019 (COVID-19) in a Golden Syrian Hamster Model: Implications for Disease Pathogenesis and Transmissibility. Clin Infect Dis. 2020 Dec 3;71(9):2428–2446. doi: 10.1093/cid/ciaa325. PMID: 32215622; PMCID: PMC7184405.

[25] Sia SF, Yan LM, Chin AWH, Fung K, Choy KT, Wong AYL, Kaewpreedee P, Perera RAPM, Poon LLM, Nicholls JM, Peiris M, Yen HL. Pathogenesis and transmission of SARS-CoV-2 in golden hamsters. Nature. 2020 Jul;583(7818):834–838. doi: 10.1038/s41586-020-2342-5. Epub 2020 May 14. PMID: 32408338; PMCID: PMC7394720.

[26] Roberts A, Vogel L, Guarner J, Hayes N, Murphy B, Zaki S, Subbarao K. Severe acute respiratory syndrome coronavirus infection of golden Syrian hamsters. J Virol. 2005 Jan;79(1):503–11. doi: 10.1128/JVI.79.1.503-511.2005. PMID: 15596843; PMCID: PMC538722.

[27] WHO SAGE Interim recommendations for use of the Pfizer-BioNtech COVID-19 vaccine, BNT162b2, under emergency use listing. WHO. 8 Jan 2021 WHO/2019-nCoV/vaccines/SAGE_recommendation/BNT162b2/2021.1

[28] FDA statement on following the authorized dosing schedules for COVID-19 vaccines. January 2021. https://www.fda.gov/news-events/press-announcements/fda-statement-following-authorized-dosing-schedules-covid-19-vaccines

[29] Kuo TY, Lin MY, Coffman RL, Campbell JD, Traquina P, Lin YJ, Liu LT, Cheng J, Wu YC, Wu CC, Tang WH, Huang CG, Tsao KC, Chen C. Development of CpG-adjuvanted stable prefusion SARS-CoV-2 spike antigen as a subunit vaccine against COVID-19. Sci Rep. 2020 Nov 18;10(1):20085. doi: 10.1038/s41598-020-77077-z. PMID: 33208827; PMCID: PMC7676267.

[30] Schnell MJ, Buonocore L, Kretzschmar E, Johnson E, Rose JK. Foreign glycoproteins expressed from recombinant vesicular stomatitis viruses are incorporated efficiently into virus particles. Proc Natl Acad Sci U S A. 1996 Oct 15;93(21):11359–65. doi: 10.1073/pnas.93.21.11359. PMID: 8876140; PMCID: PMC38062.

[31] Nègre D, Mangeot PE, Duisit G, Blanchard S, Vidalain PO, Leissner P, Winter AJ, Rabourdin-Combe C, Mehtali M, Moullier P, Darlix JL, Cosset FL. Characterization of novel safe lentiviral vectors derived from simian immunodeficiency virus (SIVmac251) that efficiently transduce mature human dendritic cells. Gene Ther. 2000 Oct;7(19):1613–23. doi: 10.1038/sj.gt.3301292. PMID: 11083469.

[32] Muñoz N,Manalastas R Jr, Pitisuttithum P, Tresukosol D, Monsonego J, Ault K, Clavel C, Luna J, Myers E, Hood S, Bautista O, Bryan J, Taddeo FJ, Esser MT, Vuocolo S, Haupt RM, Barr E, Saah A. Safety, immunogenicity, and efficacy of quadrivalent human papillomavirus (types 6, 11, 16, 18) recombinant vaccine in women aged 24-45 years: a randomised, double-blind trial. Lancet. 2009 Jun 6;373(9679):1949–57. doi: 10.1016/S0140-6736(09)60691-7. Epub 2009 Jun 1. PMID: 19493565.

[33] Keating GM, Noble S. Recombinant hepatitis B vaccine (Engerix-B): a review of its immunogenicity and protective efficacy against hepatitis B. Drugs. 2003;63(10):1021–51. doi: 10.2165/00003495-200363100-00006. PMID: 12699402.

[34] Tseng CT, Sbrana E, Iwata-Yoshikawa N, Newman PC, Garron T, Atmar RL, Peters CJ, Couch RB. Immunization with SARS coronavirus vaccines leads to pulmonary immunopathology on challenge with the SARS virus. PLoS One. 2012;7(4):e35421. doi: 10.1371/journal.pone.0035421. Epub 2012 Apr 20. Erratum in: PLoS One. 2012;7(8). doi:10.1371/annotation/2965cfae-b77d-4014-8b7b-236e01a35492. PMID: 22536382; PMCID: PMC3335060.

[35] Eichinger KM, Kosanovich JL, Gidwani SV, Zomback A, Lipp MA, Perkins TN, Oury TD, Petrovsky N, Marshall CP, Yondola MA, Empey KM. Prefusion RSV F Immunization Elicits Th2-Mediated Lung Pathology in Mice When Formulated With a Th2 (but Not a Th1/Th2-Balanced) Adjuvant Despite Complete Viral Protection. Front Immunol. 2020 Jul 29;11:1673. doi: 10.3389/fimmu.2020.01673. PMID: 32849580; PMCID: PMC7403488.

[36] Yasui F, Kai C, Kitabatake M, Inoue S, Yoneda M, Yokochi S, Kase R, Sekiguchi S, Morita K, Hishima T, Suzuki H, Karamatsu K, Yasutomi Y, Shida H, Kidokoro M, Mizuno K, Matsushima K, Kohara M. Prior immunization with severe acute respiratory syndrome (SARS)-associated coronavirus (SARS-CoV) nucleocapsid protein causes severe pneumonia in mice infected with SARS-CoV. J Immunol. 2008 Nov 1;181(9):6337–48. doi: 10.4049/jimmunol.181.9.6337. PMID: 18941225.

[37] Côté J, Garnier A, Massie B, Kamen A. Serum-free production of recombinant proteins and adenoviral vectors by 293SF-3F6 cells. Biotechnol Bioeng. 1998 Sep 5;59(5):567-75. PMID: 10099373.

